# LCLNCRdb: A Comprehensive Resource for Investigating long non-coding RNAs in Lung Cancer

**DOI:** 10.1101/2025.02.14.638263

**Authors:** Ayushi Dwivedi, Afrin Zulfia S, Mallikarjuna Thippana, Sai Nikhith Cholleti, Vaibhav Vindal

## Abstract

Lung cancer is a primary cause of death worldwide, accounting for a substantial number of mortalities. It involves several molecular mechanisms that are influenced by long non-coding RNAs (lncRNAs), a specific types of RNA molecules that do not code for proteins. Several research have revealed the importance of long non-coding RNAs (lncRNAs) in the initiation, progression, and development of resistance to lung cancer therapy. However, there are no centralized web resources or databases that collect and integrate information regarding lung cancer associated lncRNAs. This led to the development of the LCLNCRdb, a manually curated database that includes data from various sources, such as published research articles, and The Cancer Genome Atlas (TCGA) data portal. This database contains detailed information on 1102 lncRNAs that have differential expression patterns in lung cancer patients, such as lncRNA name, entrez ID, Ensemble ID, HGNC ID, NONCODE ID, lung cancer type, source, lncRNA expression pattern, experimental techniques, network analysis, and survival analysis details. The database offers a user-friendly platform for browsing, retrieving, and downloading data, and it features a dedicated submission page for researchers to share newly identified lncRNAs related to lung cancer. LCLNCRdb aims to enhance our knowledge of lncRNA deregulation in lung cancer and provides a valuable and timely resource for lncRNA research. The database is freely accessible at (https://dbtcmi.in/tools/lclncrdb/main.html).

## 1. Introduction

Lung cancer is a major health concern, causing millions of deaths annually. It is the second most prevalent form of cancer and the leading cause of cancer-related deaths. Factors, such as smoking, genetics, and exposure to harmful substances, contribute to its development. Non-small cell lung cancer (NSCLC) is the most prevalent type, representing 85% of all instances, and is categorized into the adenocarcinoma (LUAD) and squamous cell carcinoma (LUSC) subtypes. Long non-coding RNAs (lncRNAs) play crucial roles in the regulation of gene expression across multiple levels, including epigenetic, transcriptional, and post-transcriptional processes. They can affect chromosome structure, recruit chromatin-modifying enzymes, and interact with transcription factors to either enhance or repress gene expression [1–4]. Additionally, they can affect mRNA stability and translation and function as competing endogenous RNAs (ceRNAs) that sequester microRNAs (miRNAs) [5]. Dysregulation of lncRNAs can contribute to abnormal gene expression patterns, potentially leading to the development of various diseases, including cancer [6].

Long non-coding RNAs (lncRNAs) contributes to the cancer progression, and various databases have addressed diverse aspects of lung cancer progression. However, none have specifically focused on lncRNAs associated with lung cancer progression. In lung cancer, dysregulated lncRNAs promote cell growth, migration, and invasion, and inhibit apoptosis [7]. For example, lncRNA DANCR is upregulated in various cancers and is associated with increased cell proliferation and invasion. Similarly, MALAT1 and H19 have been implicated in lung cancer development. The dysregulation of lncRNAs can also affect key signaling pathways and regulatory networks, such as the p53 tumor suppressor pathway [6]. Therefore, lncRNAs are important regulators of gene expression, and their dysregulation can lead to the disruption of cellular homeostasis and lung cancer progression. The development of a database for long non-coding RNAs (lncRNAs) associated with lung cancer is underscored by a growing body of evidence highlighting their significant roles in cancer biology. They participate in the regulation of gene expression at various levels and are associated with the initiation, progression, and prognosis of lung cancer [8–9]. Furthermore, long non-coding RNAs (lncRNAs) have emerged as promising candidates for use as diagnostic and prognostic biomarkers, as well as potential targets for therapeutic intervention [10–11]. This knowledge gap presents a compelling case for a dedicated database that could consolidate current and future research findings, facilitating a more comprehensive understanding of lncRNA functions and interactions in lung cancer.

Several databases exist that compile and organizes data associated with lncRNAs and cancer. Among these, Lnc2Cancer is a manually curated database that contains detailed mechanisms of lncRNA regulation in cancer [12–13]. It contains comprehensive information on the mechanisms by which lncRNAs regulate cancer development. Another database, CRlncRNA, was created by Wang and focuses on the functional roles of cancer-related lncRNAs [14]. In addition, it provides information on the clinical and molecular characteristics of these lncRNAs. In addition, LncRNADisease offers information about lncRNA-disease associations, along with details on transcriptional regulatory relationships and a confidence score for each association [15]. These databases are invaluable resources for researchers and clinicians seeking to understand and explore the roles of lncRNAs in cancer. Nevertheless, none of these investigations have specifically addressed long non-coding RNAs (lncRNAs) linked to lung cancer, which represents a substantial gap in the current body of knowledge.

To address the lack of information on long non-coding RNAs (lncRNAs) associated lung cancer, a comprehensive database designated as the LCLNCRdb was developed. This database includes 1102 differentially expressed lncRNAs in lung cancer, which were manually curated from both the literature and TCGA databases. The LCLNCRdb offers a range of features, such as lncRNA expression patterns, target information, type of lung cancer, source data, experimental techniques, survival analysis, and network analysis. The database is user friendly, allowing users to easily browse, retrieve, and download data. In addition, users can submit newly validated lncRNAs related to lung cancer. The LCLNCRdb was developed using the XAMPP webserver, HTML, PHP 8.2.0, JavaScript, MySQL, Bootstrap 5, and DataTables plug-in. The database is freely accessible at https://dbtcmi.in/tools/lclncrdb/main.html and is a valuable resource for researchers studying lung cancer and lncRNAs, with the potential to greatly advance research in this field.

## 2. Web resource content and methods

The data for transcriptomic profiling of LUAD and LUSC tumor and normal samples were obtained using the TCGAbiolinks R package [16] from the TCGA-GDC portal. The projects were TCGA-LUAD and TCGA-LUSC, comprising 598 and 551 samples, respectively. Of these, 537 LUAD and 502 LUSC samples were tumor-positive, whereas 59 LUAD and 49 LUSC samples were normal. Following pre-processing to remove duplicates and low read count entries, gene symbols were mapped to coding and non-coding entities to classify genes as mRNA, lncRNAs, and miRNAs. Apart from this, a comprehensive search of the PubMed database was conducted to identify lncRNAs related to lung cancer using the entrez_search function of the rentrez R package [17] up until Jan 24, 2024. Keywords such as "long non-coding RNA, " "lncRNA, " "long non-coding, " and "lung cancer" were used to retrieve the information. The search yielded 1924 hits, reporting lncRNAs associated with lung cancer development, progression, diagnosis, and treatment (Figure 1). The information on lncRNAs was obtained from the HGNC database, and their sequences were sourced from the Ensembl database. The Gene Cards database was used to determine their association with lung cancer and other forms of cancer. This information was systematically stored and managed using the MySql data tables.

**Figure 1:**
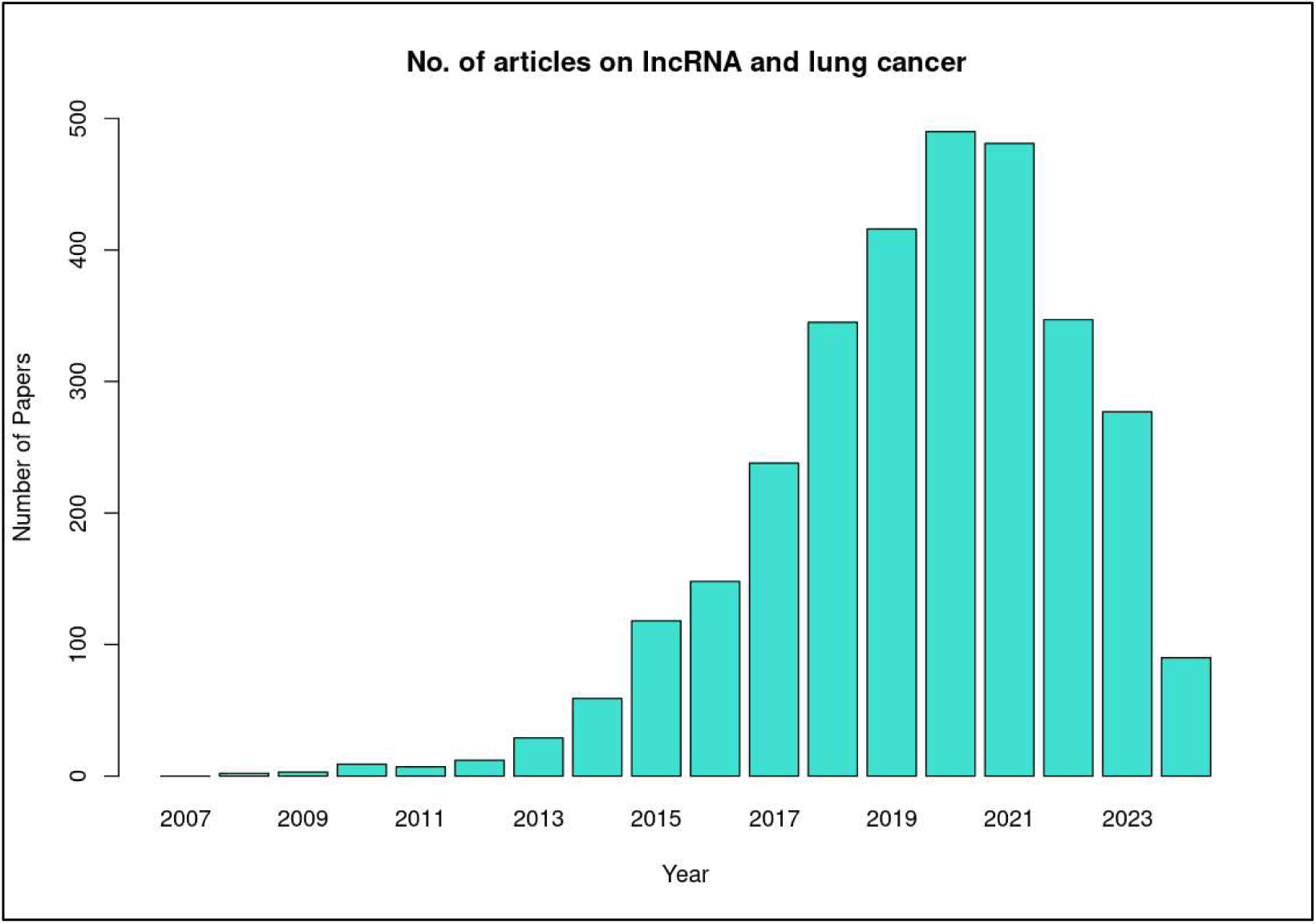
No. of articles on lncRNA and Lung cancer

## 3. Analysis of transcriptomic data

### 3.1 Differential gene expression analysis

A differential gene expression analysis was carried out using the DESeq2 package [18], which allowed us to identify genes with significant changes in expression. The filtering criteria for differentially expressed genes were set at log2FC =< 2 and a adjusted P value of less than 0.05, which were used to screen out genes that exhibited significant changes in expression. Differentially expressed genes were then categorized into protein-coding and non-coding groups. Additionally, the regulatory networks of the target genes were examined using differentially expressed non-coding elements such as DE-miRNA (DEM) and DE-lncRNA (DEL).

### 3.2 Target prediction and network construction

#### 3.2.1 Prediction of target miRNAs of DE-lncRNA

In this study, 619 and 931 lncRNAs were identified as differentially expressed in LUAD and LUSC, respectively, of which 448 were common between the two cancer types, resulting in a total of 1, 102 unique DE-lncRNAs. To identify their target miRNAs, two databases were utilized: miRcode [19], which contains 10, 000 long non-coding RNAs, and lncRNASNP2 [20], which encompasses experimentally validated microRNA-long non-coding RNA interactions.

#### 3.2.2 Prediction of target mRNAs of DE-miRNA

Our study utilized the miRDB [21] database to identify mRNAs that are regulated by DE-miRNAs. This database comprises an extensive collection of predicted miRNA-mRNA interactions. The investigation primarily focused on mRNAs that exhibited differential expression in the target genes of DE-miRNAs to elucidate specific interactions.

#### 3.2.3 Target-gene interaction network construction

To construct the lncRNA-miRNA-mRNA competing endogenous RNA (ceRNA) network, DE-lncRNA-DE-miRNA and DE-miRNA-DE-mRNA target networks were integrated for both LUAD and LUSC. This approach facilitates the understanding of complex regulatory mechanisms by mapping the interactions between different RNA molecules in lung cancer subtypes. Specifically, there were 5 such DE-miRNAs for LUAD and 6 for LUSC. Consequently, only 5 DE-miRNAs and their corresponding target DE-mRNAs were included from LUAD and 6 DE-miRNAs and their corresponding target DE-mRNAs from LUSC in constructing the network.

### 3.3 Survival Analysis

Survival analysis was carried out to investigate the effect of differentially expressed genes, lncRNAs, and miRNAs on patient survival in the LUAD and LUSC groups. The lncExplore database [22] was used to generate survival curves for patients with high or low expression levels of DE-lncRNAs over time. The Kaplan-Meier plotter [23] tool was used to assess the prognostic performance of DE-miRNAs and DE-mRNAs from the ceRNA network, by calculating the Cox proportional hazards ratio (HR) > 1 and log-rank p-value cutoff < 0.05 to identify poorly prognosed genes. Finally, the survival probability of patients with low or high expression levels of DE-miRNAs and DE-mRNAs over time (in months) was compared.

### 3.4 Database Construction

The database was constructed using the XAMPP web server, HTML, PHP 8.2.0, and JavaScript for the front-end; MySQL for the backend to store database tables; and Bootstrap 5 for styling purposes. Additionally, the DataTables plugin was employed to present tables with a large amount of data in an organized manner.

## 4. Results

### 4.1 Data summary

The LCLNCRdb is a comprehensive database that contains information on 1, 102 long non-coding RNAs (lncRNAs) that exhibit differential expression patterns in lung cancer patients. In addition to basic information, the database includes data on target-gene regulatory interactions and predicted prognostic performance in patients with lung cancer.

### 4.2 Web interface and usage

#### 4.2.1 User interface modules

The primary goal of the LCLNCRdb is to identify and analyze long non-coding RNAs (lncRNAs) that exhibit differential expression in lung cancer. They have the potential to significantly contribute to lung cancer development and progression and may serve as biomarkers or therapeutic targets. The database provides information on 1, 102 lncRNAs and their expression levels in lung adenocarcinoma and squamous cell carcinoma. LCLNCRdb offers a user-friendly interface for exploring differentially expressed lncRNAs, target networks, and survival analysis plots, and users can download the information in csv format from the download module. The platform provides four user-friendly web interfaces to access the database.

#### 4.2.2 Homepage

Our database comprises 1102 long non-coding RNAs (lncRNAs) that exhibit differential expression profiles along with their corresponding target networks and survival analysis plots. The homepage of our database, LCLNCRdb, features a fixed navigation bar at the top and displays the name of the database, as shown in Figure 2.

**Figure 2:**
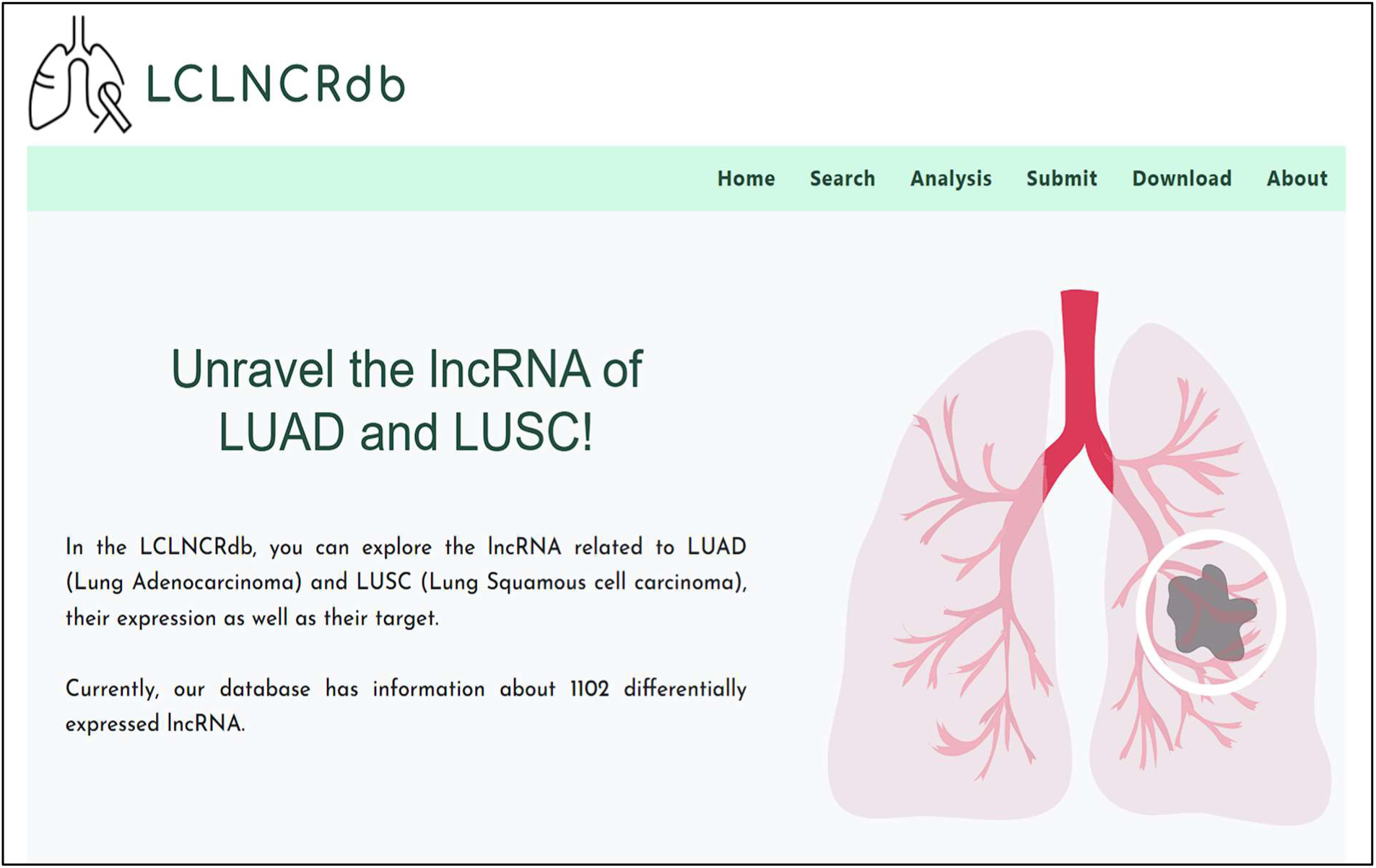
Homepage of LCLNCRdb

#### 4.2.3 Navigation Tabs

The database’s user interface includes a navigation bar with six sections: Home, Search, Analysis, Submit, Download, and About. The Search tab allows users to search for long non-coding RNA (lncRNAs) using gene names, sequence information, or a list of lncRNAs. The Analysis tab offers a network analysis of target genes and competing endogenous RNA (ceRNA) networks, as well as survival analysis plots. The About tab provides information on how to use the database and includes contact details, as illustrated in Figure 3.

**Figure 3(a):**
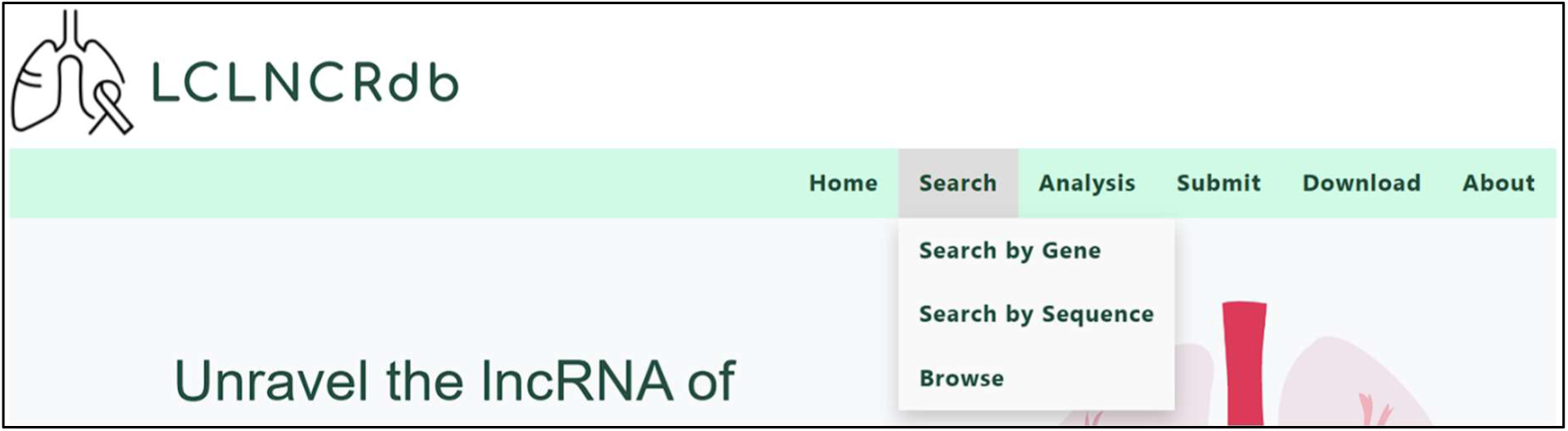
Search module option of LCLNCRdb

**Figure 3(b):**
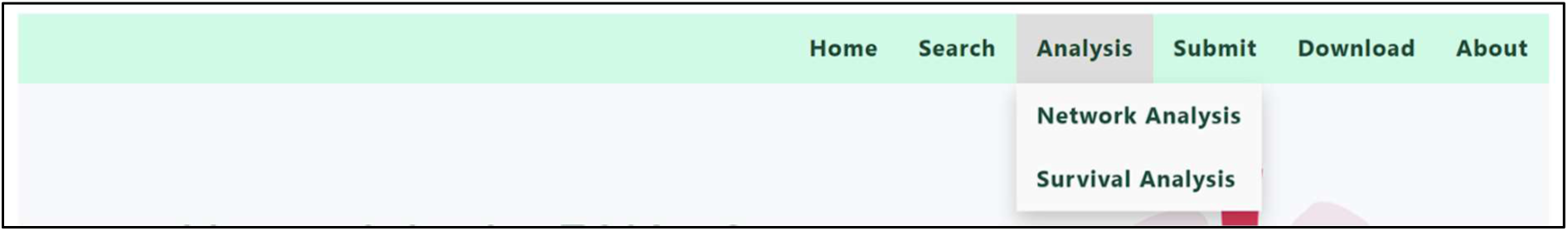
Analysis module option of LCLNCRdb

**Figure 3(c):**
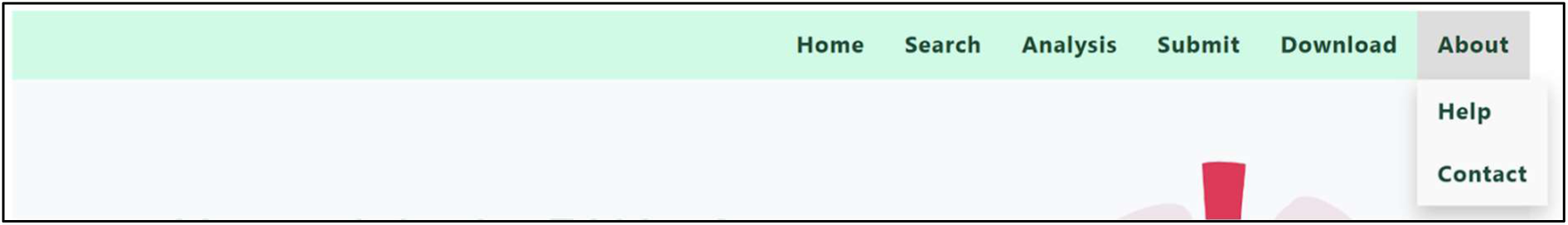
About module option of LCLNCRdb

#### 4.2.4 Search module

The search module features three distinct search tabs that can be accessed via a drop-down menu: search by gene, search by sequence, and browsing. The user has the option to select from the two search methods. They can search for a gene by entering its official symbol, Entrez ID, HGNC ID, Ensemble ID, NONCODE ID or by submitting the FASTA sequence of the gene.

The following information is summarized in Figure 4, which provides a detailed breakdown of each long non-coding RNA (lncRNA): The figure displays the lncRNA’s unique identifier (ids), aliases, map location, source, types, and expression pattern. By clicking on a lncRNA’s RefSeq ID, the user can access its FASTA sequence and view the network of microRNAs (miRNAs) targeted by the lncRNA below the sequence. Additionally, the survival plot and p-value shown in the figure indicate the prognostic significance of the lncRNAs.

**Figure 4:**
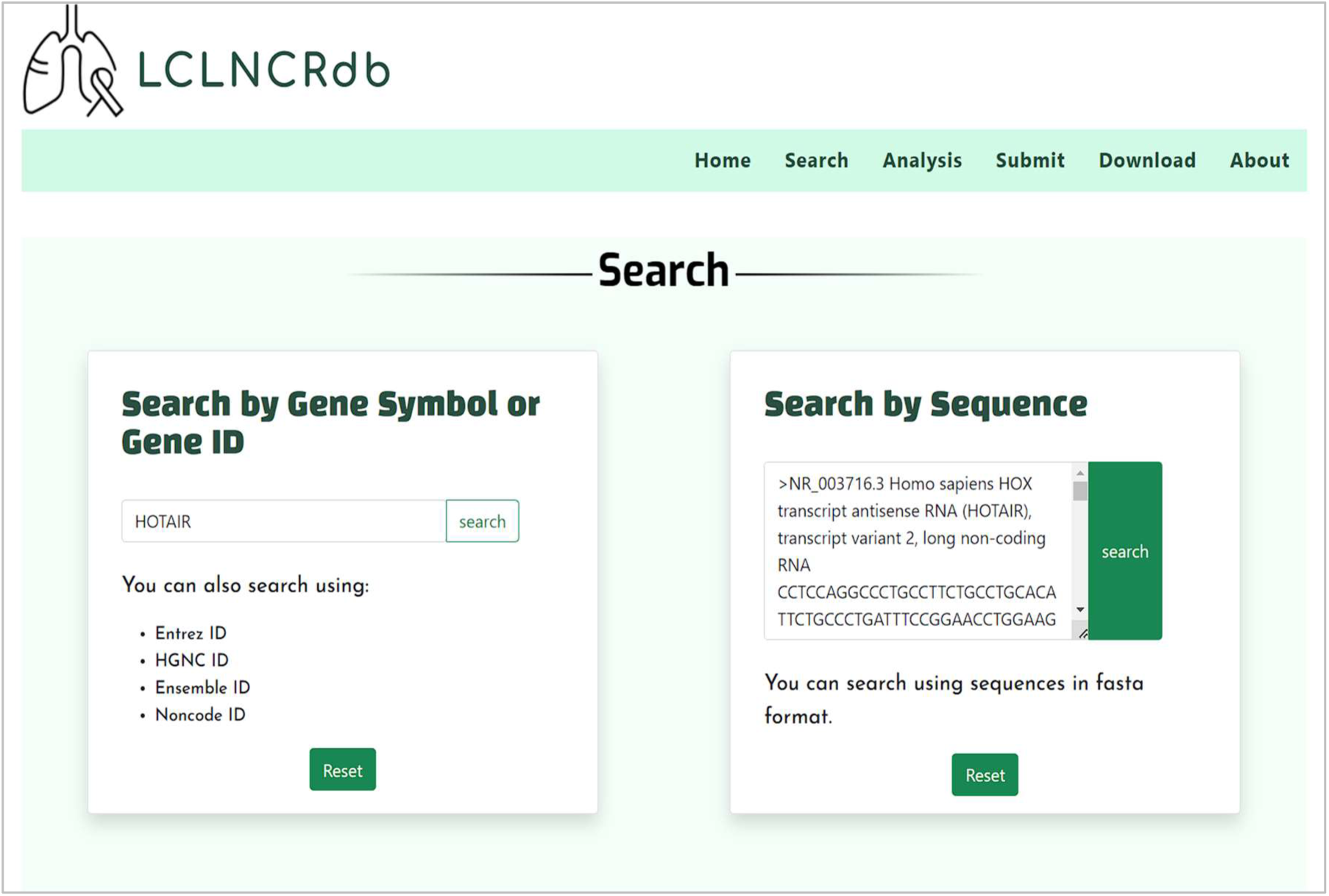
Search page of LCLNCRdb

**Figure 4:**
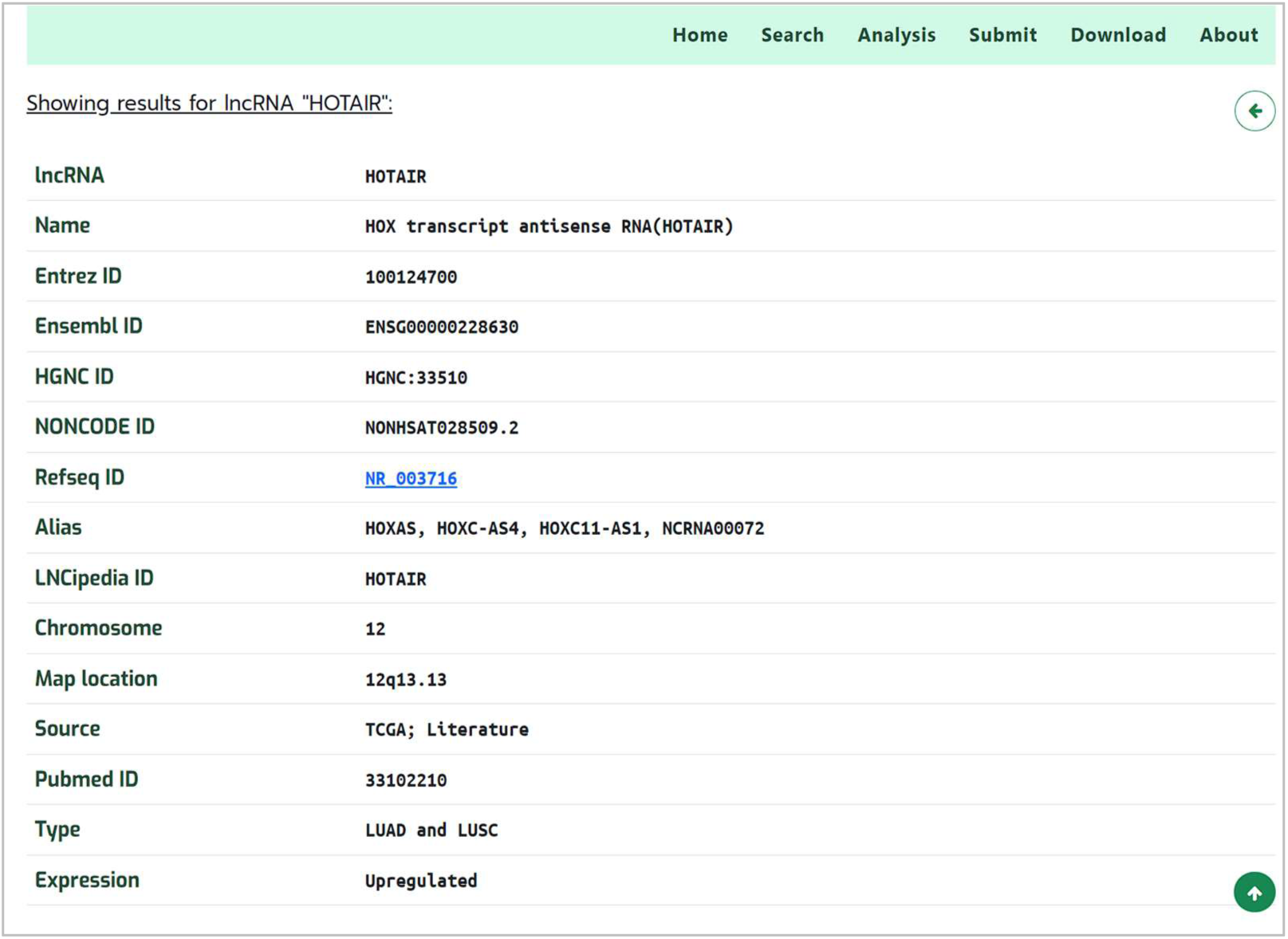
Showing search results (a) General details of lncRNA “HOTAIR”

**Figure 4:**
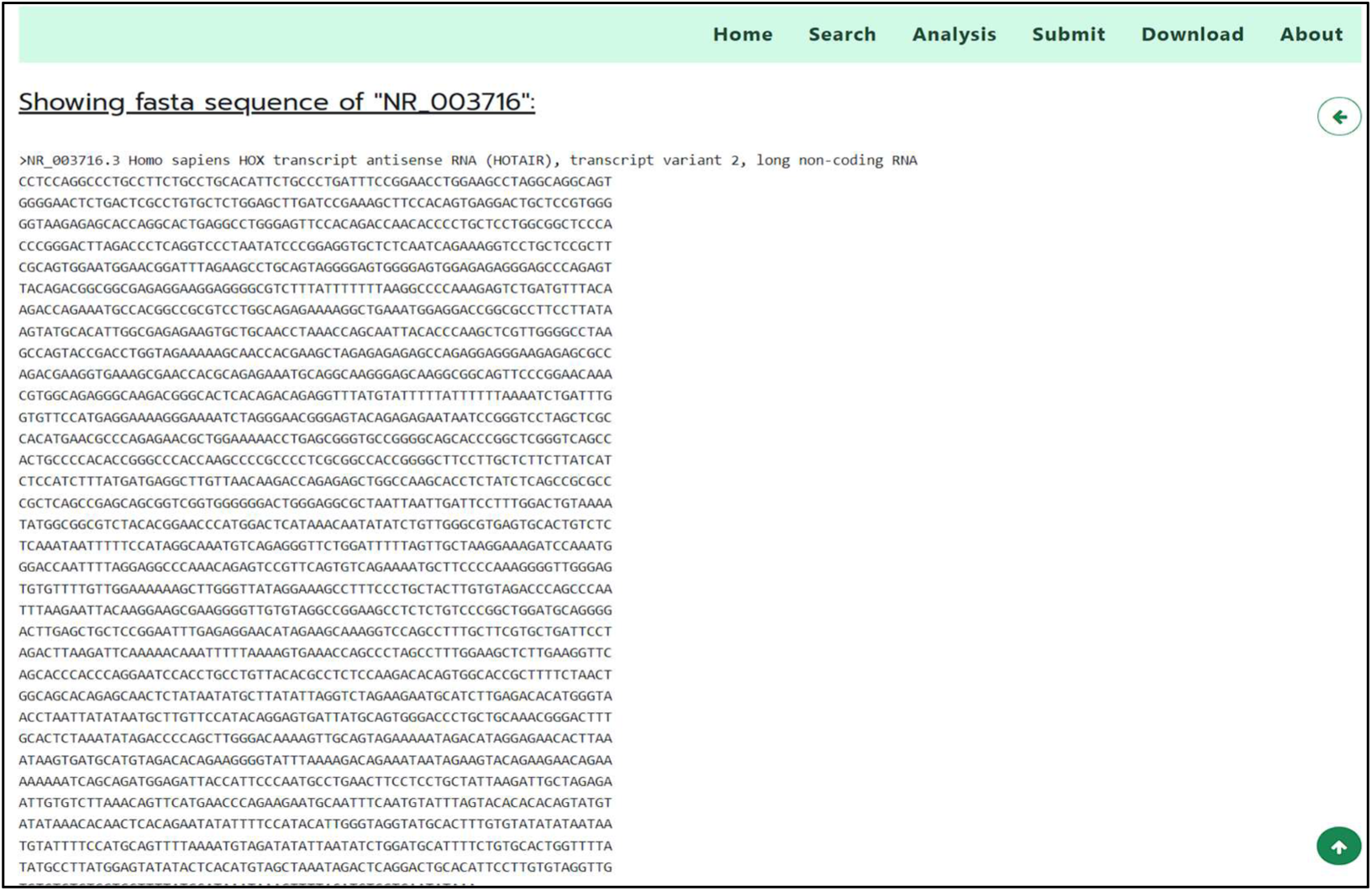
Showing search results (b) RefSeq fasta sequence of HOTAIR

**Figure 4:**
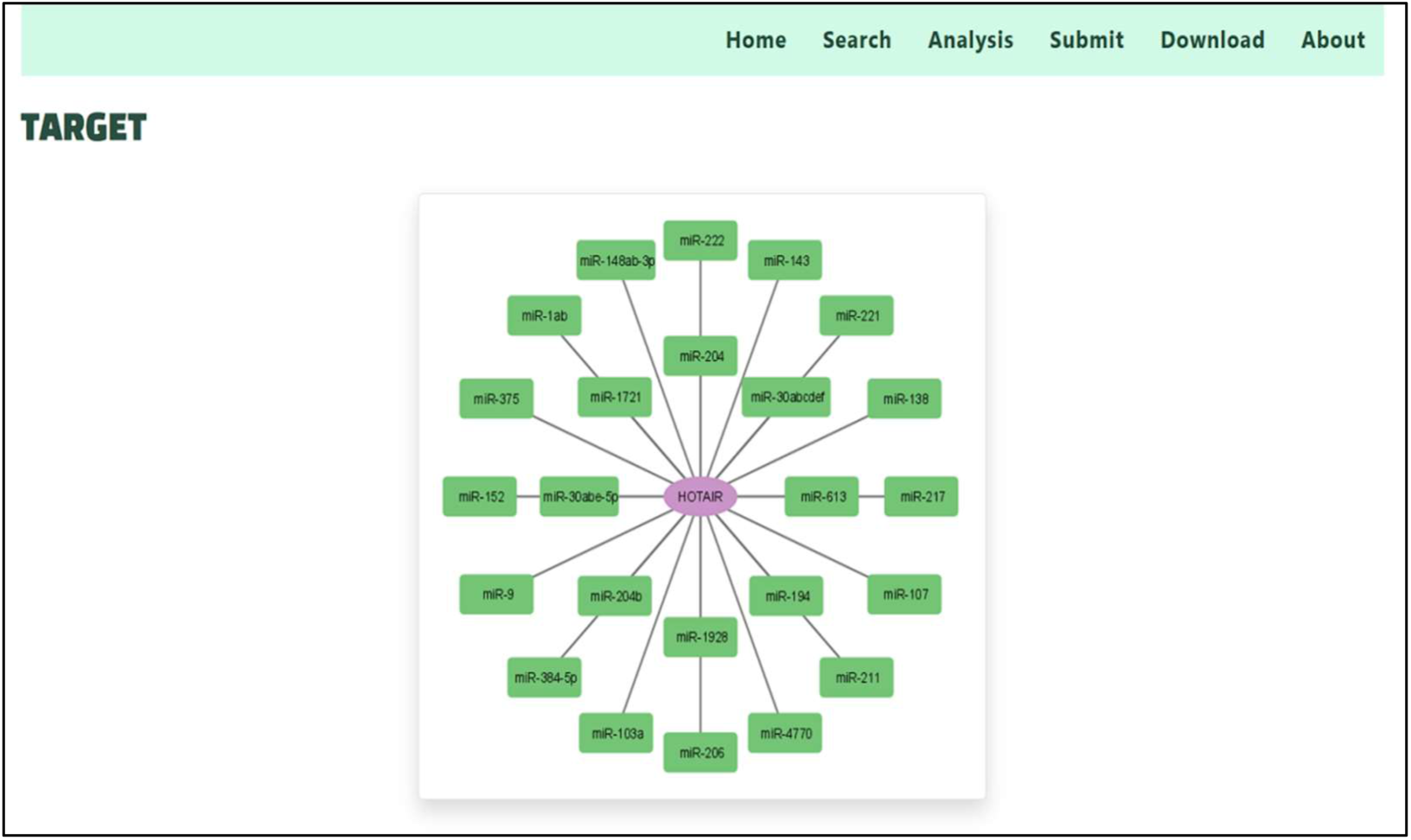
Showing search results (c) miRNA - target network of HOTAIR

**Figure 4:**
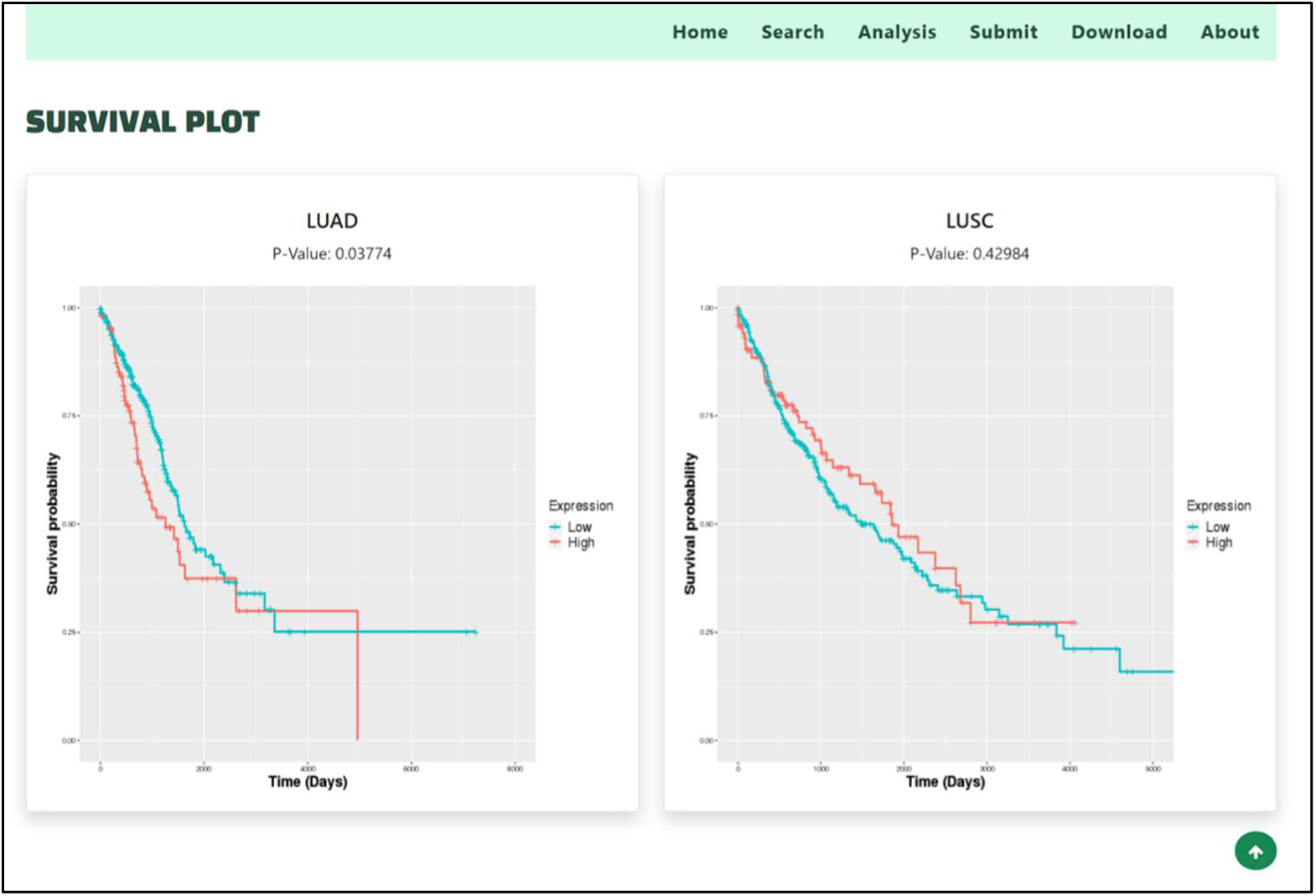
Showing search results (d) Survival plot of HOTAIR in LUAD and LUSC

#### 4.2.5 Browse module

The section for browsing the table displays all 1102 long non-coding RNAs (lncRNAs) with columns for their classification, expression pattern, and origin. Users can refine their search by selecting specific categories, such as lung adenocarcinoma (LUAD), lung squamous cell carcinoma (LUSC), or those that are shared by both. Additionally, data can be filtered based on the expression pattern of lncRNAs, which can be upregulated, downregulated, or both. Furthermore, the table can be arranged in ascending or descending order, as shown in Figure 5(a) and 5(b).

**Figure 5:(a).**
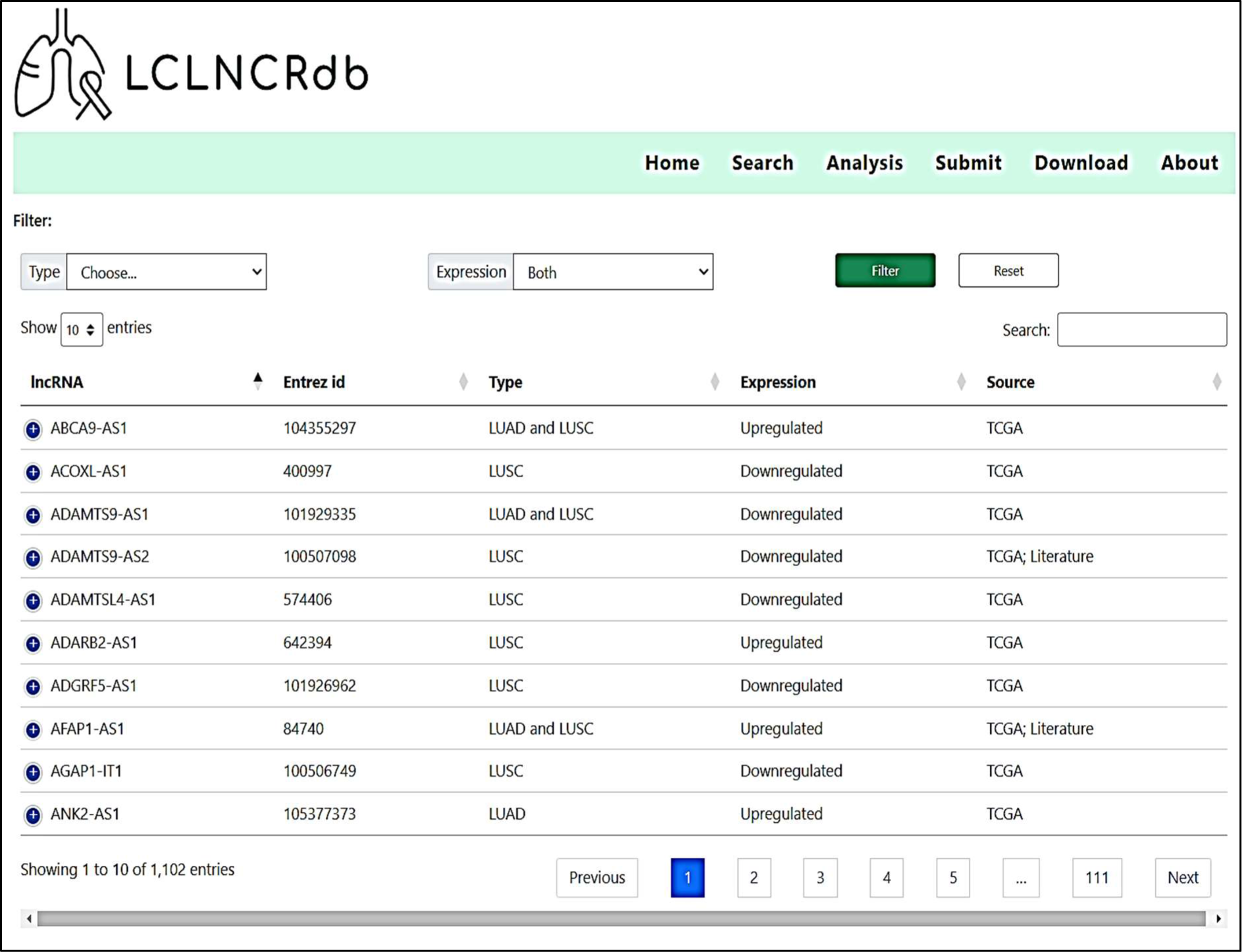
Browse page showing all 1102 lncRNA

**Figure 5:(b).**
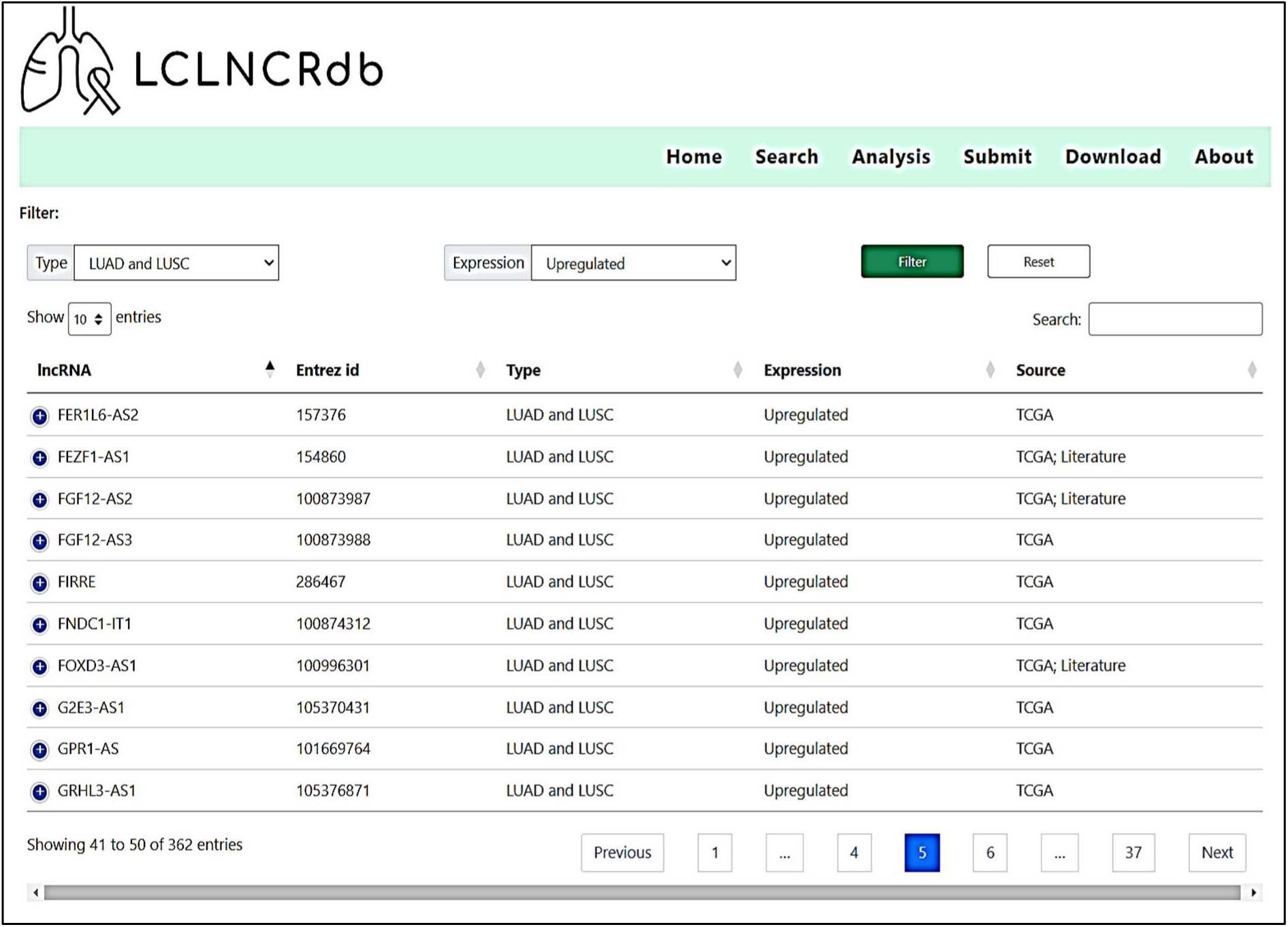
Browse page LUAD and LUSC type filter and Upregulated expression filter

#### 4.2.6 Analysis module

The Analysis tab offers two distinct modules: network and survival analyses.

##### 4.2.6.1 Network Analysis module

The Network analysis module offers three distinct types of target networks: DElncRNA-DEmiRNA, DElncRNA-DEmiRNA-DEmRNA, and Competitive endogenous RNA (ceRNA) networks, which are specifically designed to analyze miRNAs in LUAD and LUSC. To visualize these networks and offer interactive features, the Cytoscape.js JavaScript library version 3.2.4 was used. The table below the network displays the centrality measures of each gene sorted by degree, in descending order, as depicted in Figure 6(a) and 6(b).

**Figure 6:(a).**
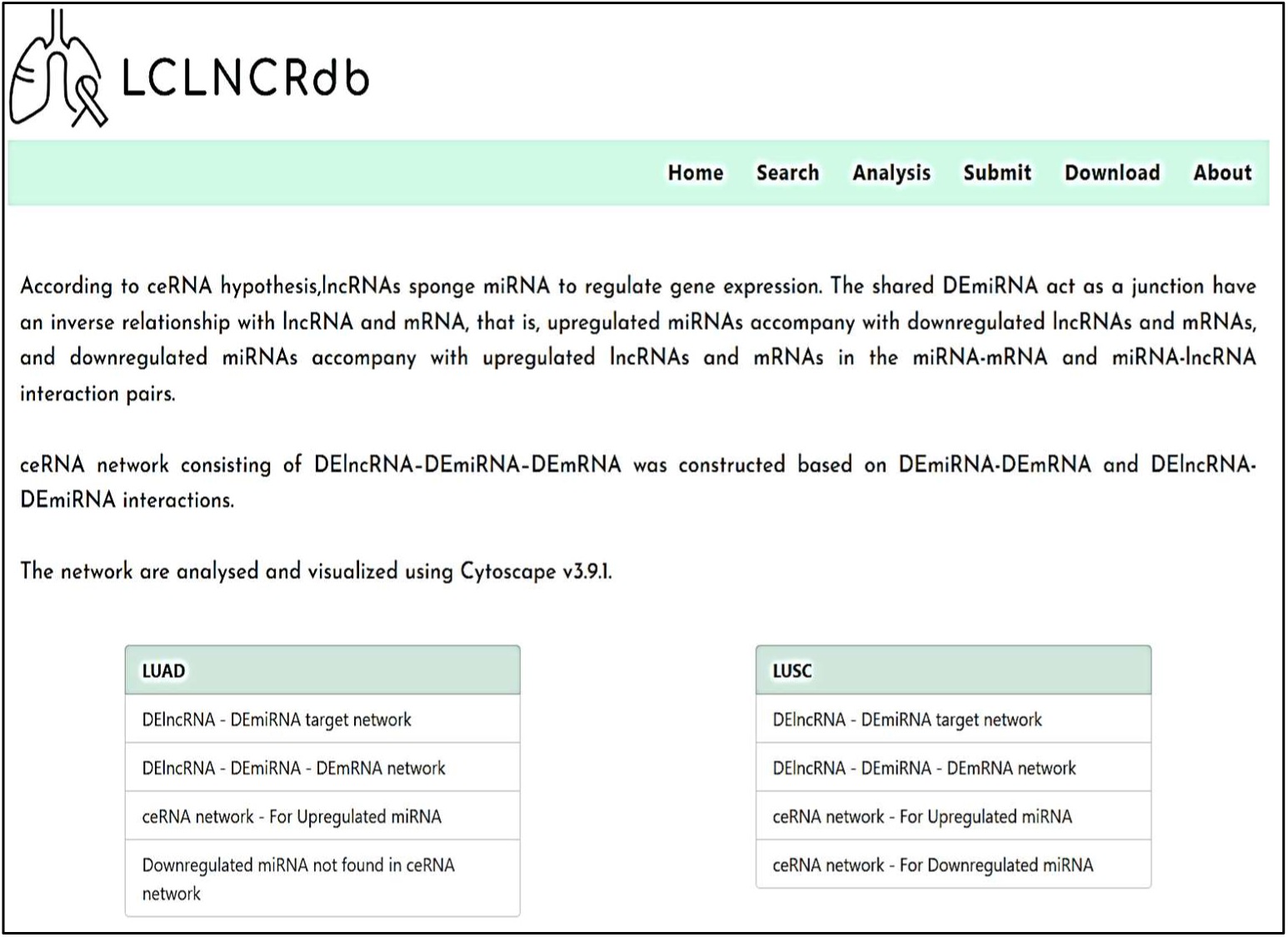
Network analysis page showing the networks available

**Figure 6:(b).**
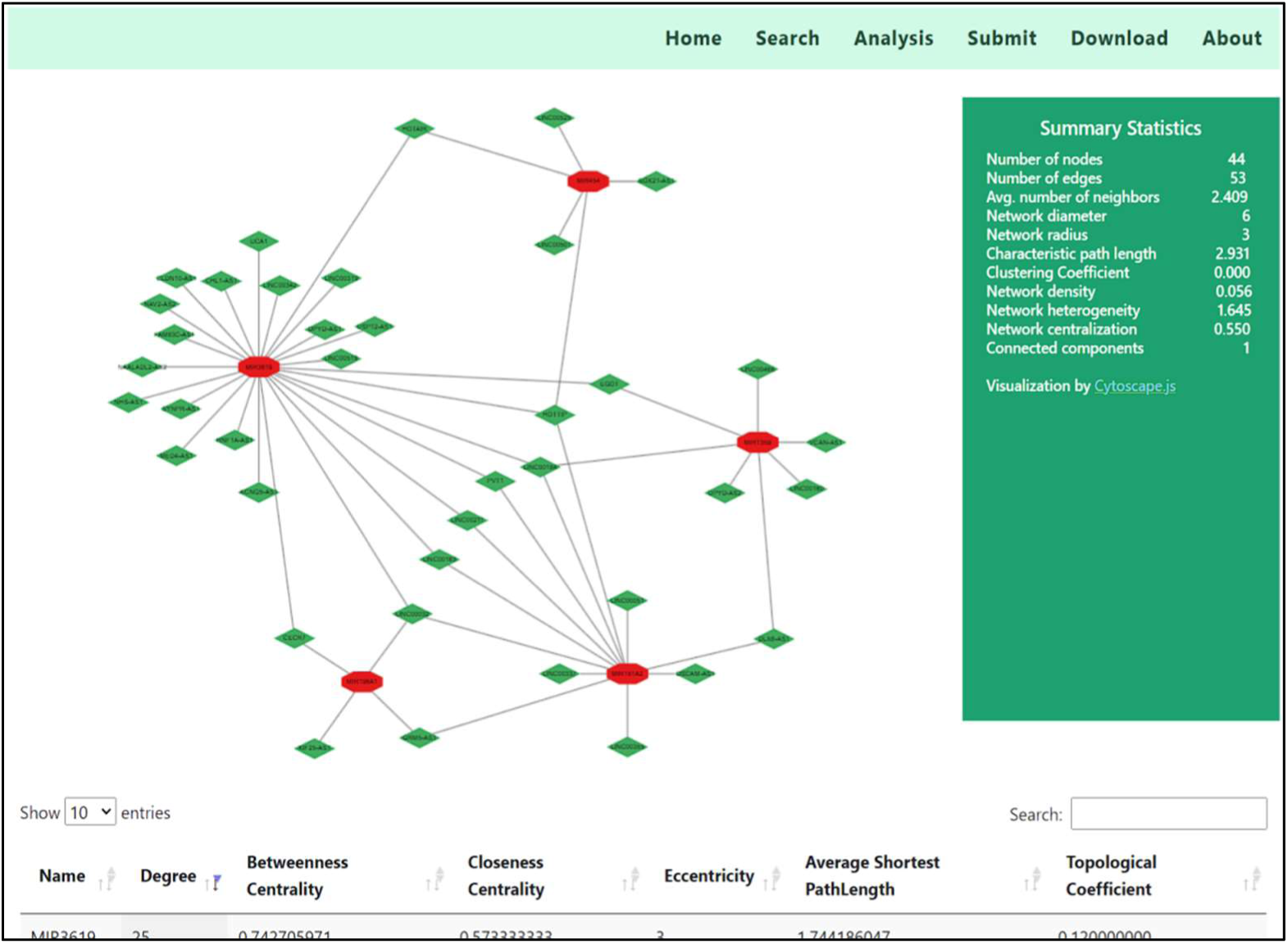
Display of DE-lncRNA – DE-miRNA target network of LUAD

##### 4.2.6.2 Survival Analysis module

The survival analysis module enables researchers to examine overall survival based on long non-coding RNA (lncRNA) expression utilizing the Kaplan-Meier survival plot. To assess the survival of a given gene, researchers input the lncRNA name into the designated field and initiate the search function, which subsequently generates the survival plot for that specific gene. The resulting plot displays the lncRNA plot along with the messenger RNA (mRNA) or microRNA (miRNA) associated with the competing endogenous RNA (ceRNA) network of lung adenocarcinoma (LUAD) and lung squamous cell carcinoma (LUSC). The plot presents both LUAD and LUSC if the gene/lncRNA is shared by both types or only one type if it is exclusive to one type, as illustrated in Figure 7(a), 7(b), and 7(c).

**Figure 7(a):**
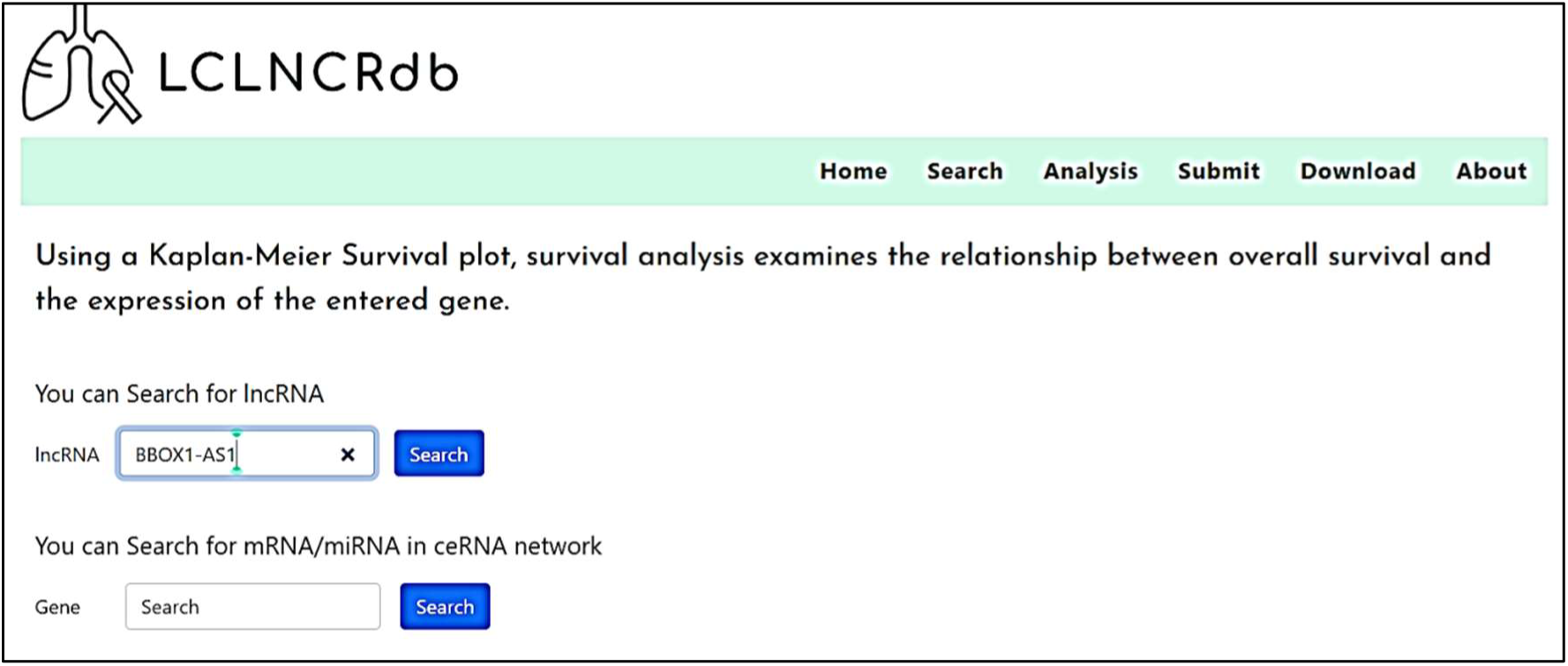
Survival Analysis page

**Figure 7(b):**
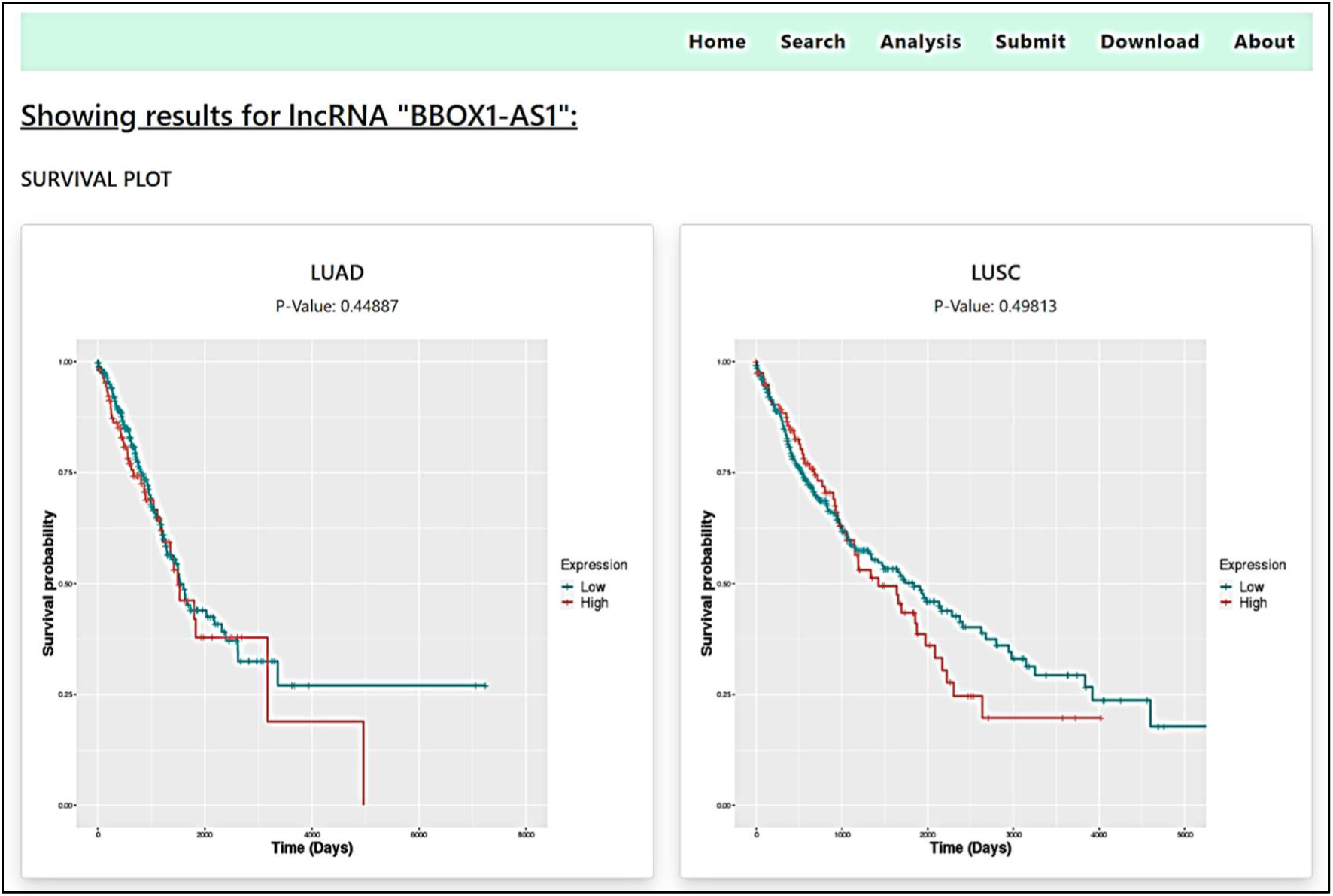
Representation of Kaplan-Meier survival plot for the lncRNA “BBOX1-AS1”

**Figure 7(c):**
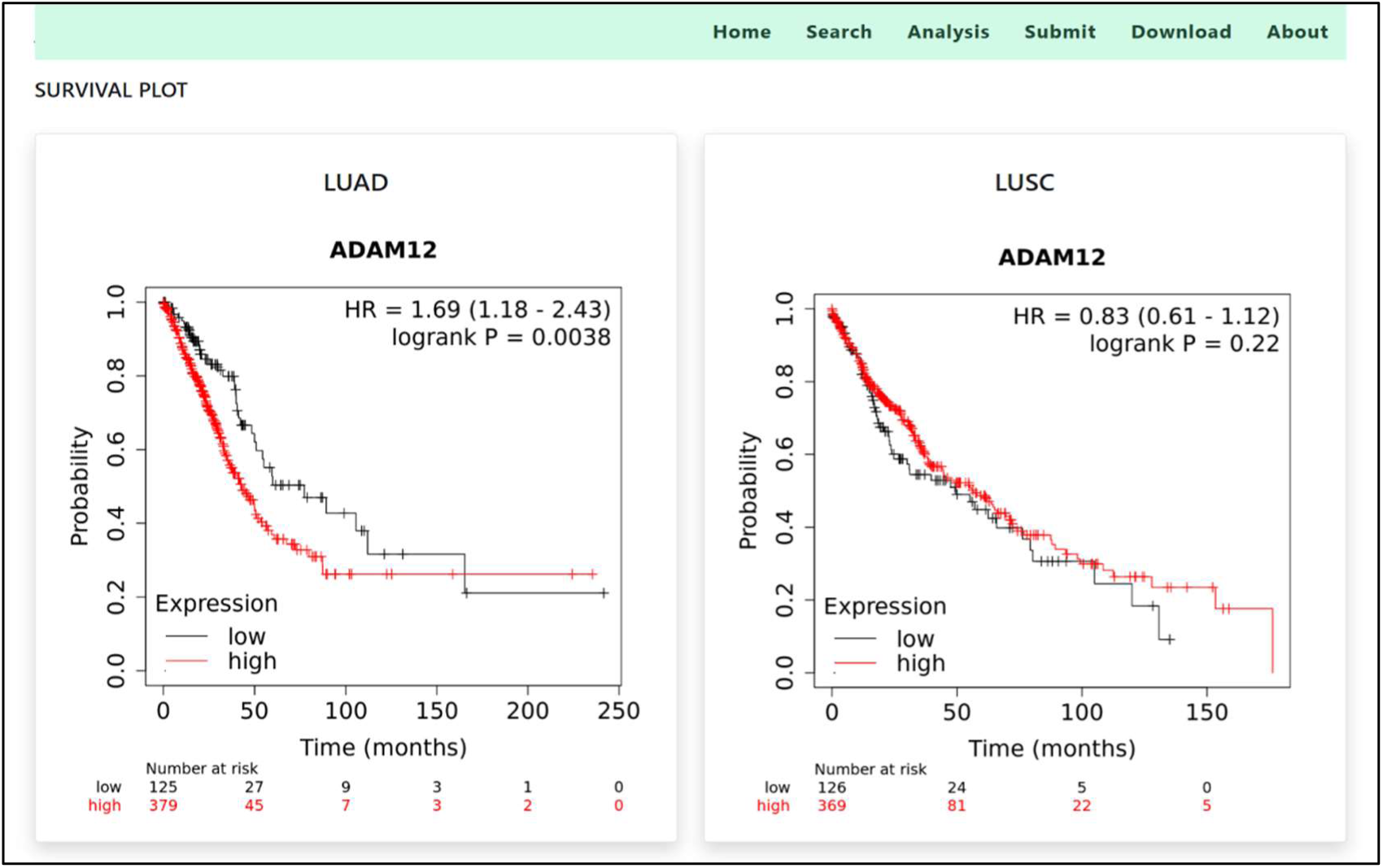
Representation of Kaplan – Meier survival plot for the mRNA “ADAM12”

#### 4.2.7 Submit module

The procedures depicted in Figure 8(a) and 8(b) are essential for effectively completing the Submit page form. To guarantee the successful submission of user enquiry or fresh information pertaining to lncRNAs and lung cancer, it is essential to verify that all fields are correctly completed. This includes the name, email address, subject, and message. Subsequently, click the Submit button to transmit the form. Upon validation, the form will be sent as an email.

**Figure 8(a):**
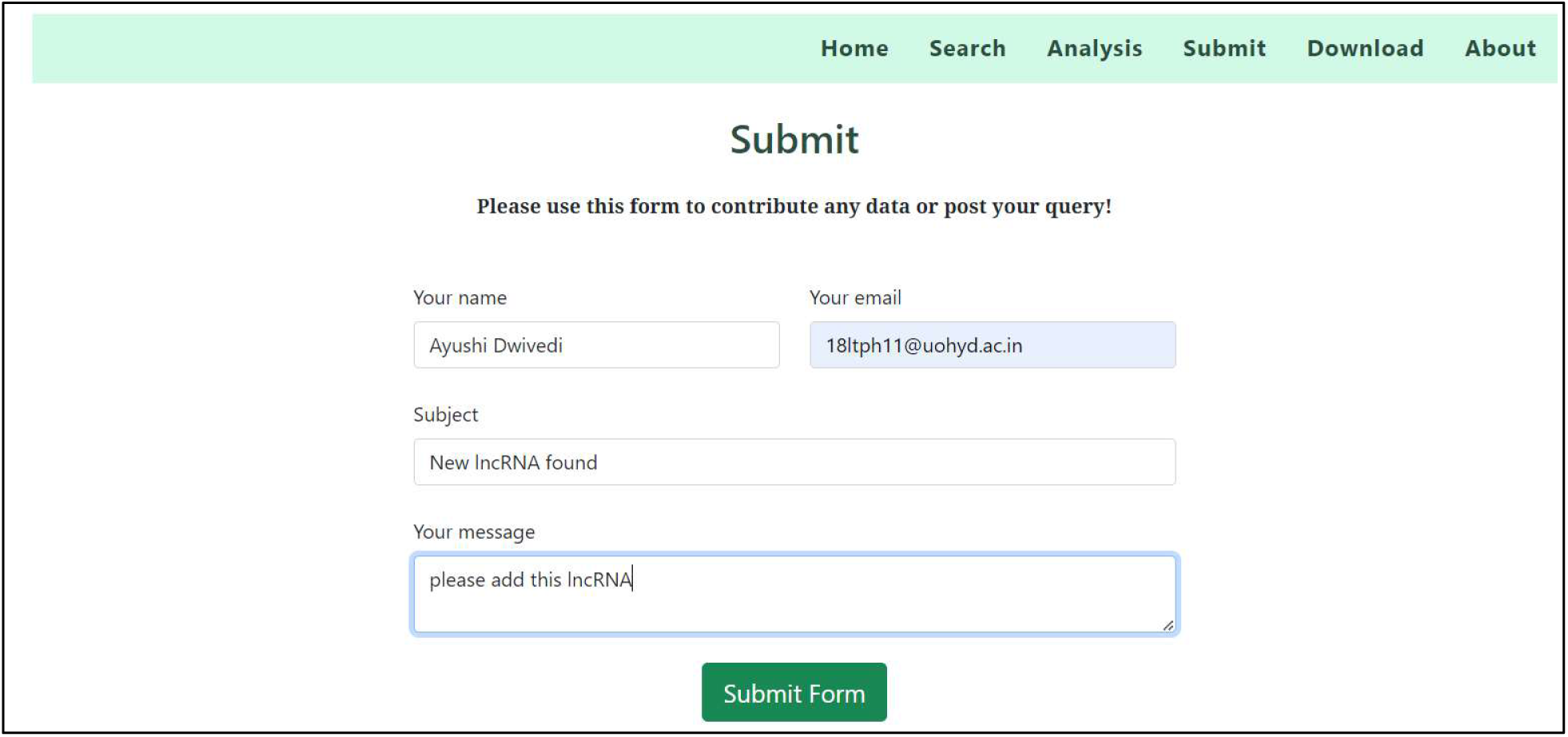
Submit page

**Figure 8:(b).**
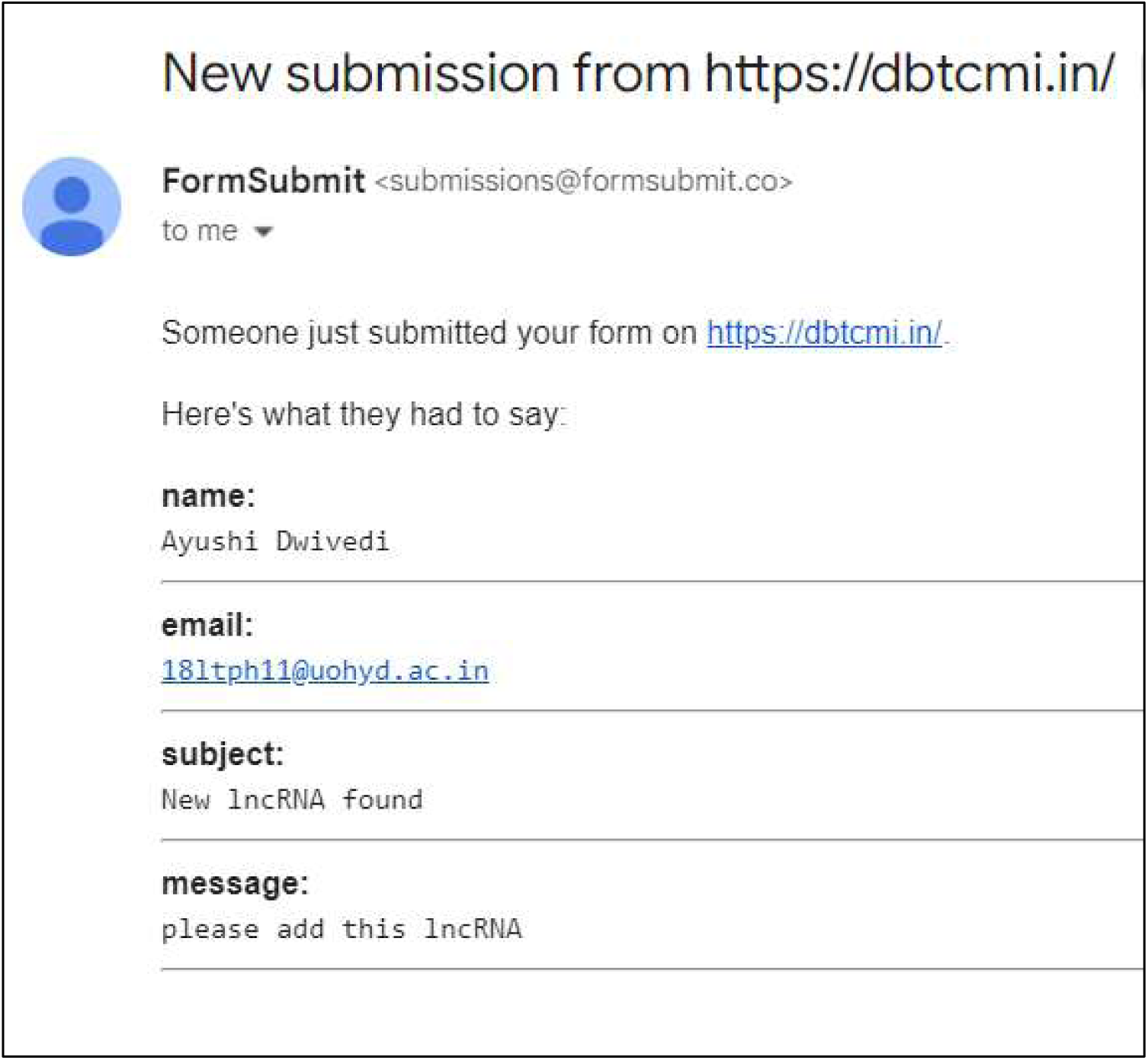
SubmiLed form as an email

#### 4.2.8 Download module

The 1102 lncRNAs in LUAD and LUSC, as well as their shared lncRNAs, can be accessed on the download page, which also includes network analysis and centrality measures. The data can be downloaded in the CSV format, as shown in Figure 9.

**Figure 9:**
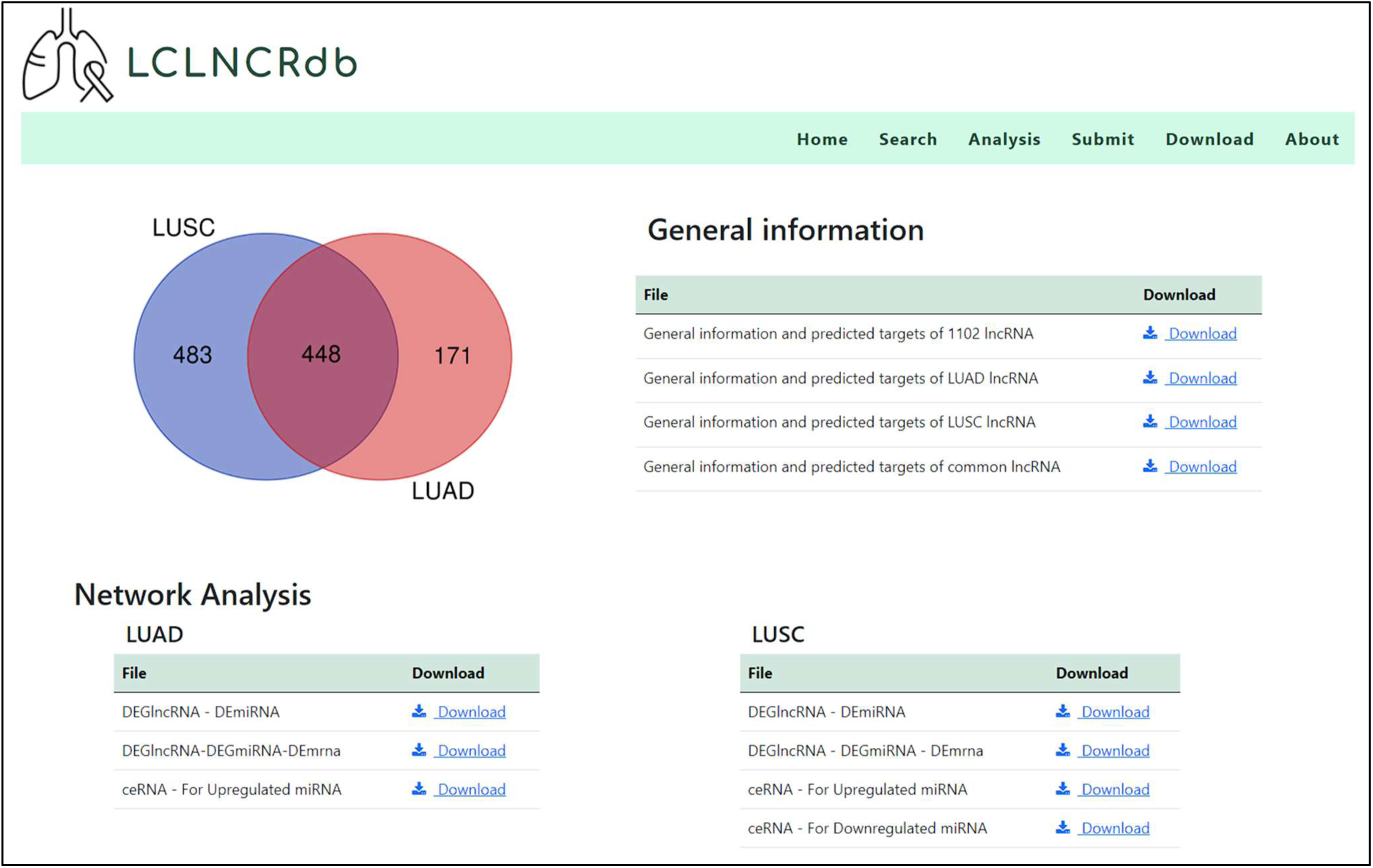
Download page of LCLNCRdb

## 5. Summary and future directions

### Summary

Lung cancer, a major global health concern, is intricately associated with the dysregulation of long non-coding RNAs (lncRNAs), which play a crucial role in the disease’s onset, progression, and treatment resistance. Despite their significance, there is a notable absence of centralized databases that compile comprehensive data on lncRNAs related to lung cancer. This gap in resources hinders the consolidation of crucial information that could potentially advance research and therapy development in this field. To address this gap, the LCLNCRdb database was developed, which includes data from various sources, such as published research articles, and TCGA. The database contains detailed information on 1102 lncRNAs that have differential expression in lung cancer patients, including lncRNA name, entrez ID, Ensemble ID, HGNC ID, NONCODE ID, lung cancer type, source, lncRNA expression pattern, experimental techniques, network analysis, and survival analysis details. LCLNCRdb offers a user-friendly interface that enables users to browse, retrieve, and download data, as well as a dedicated submission page for researchers to share newly indentified long non-coding RNAs (lncRNAs) related to lung cancer. LCLNCRdb aims to enhance our understanding of lncRNA deregulation in lung cancer and provides a valuable and timely resource for lncRNA research. The database is freely accessible at https://dbtcmi.in/tools/lclncrdb/main.html

LCLNCRdb is a comprehensive and user-friendly database that compiles information on differentially expressed long non-coding RNAs (lncRNAs) and their targets related to lung cancer, sourced from various data sources, providing an accessible and thorough resource for researchers. This resource provides extensive details regarding lncRNAs associated with lung cancer, including interactive and survival plots, to aid downstream analyses, such as identifying biomarkers and selecting target genes for future experiments. LCLNCRdb contains information on 1102 lncRNAs related to lung cancer and freely accessible. Users can retrieve different types of target networks, such as DE-lncRNA-DE-miRNA, DE-lncRNA-DEmiRNA-DE-mRNA, and competitive endogenous regulatory networks for both upregulated and downregulated miRNAs. The database aims to curate scientific literature on lncRNAs associated with lung cancer with high quality and efficiency, ensuring that the information is relevant, comprehensive, and up-to-date. LCLNCRdb also strives to improve the user experience and add new relevant content. To our knowledge, there are no other specific and up-to-date databases dedicated to lncRNAs associated with lung cancer. This database, which is specifically designed for lung cancer and its associated lncRNAs, provides a comprehensive resource for researchers. The study explored the functions and underlying processes of long non-coding RNAs (lncRNAs) in lung cancer, aiming to identify potential biomarkers and therapeutic targets.

## Acknowledgment

The authors gratefully acknowledge the DBT-Centre for Microbial Informatics (https://dbtcmi.in) for hosting the LCLNCRdb database. Vindal V would like to acknowledge the Institution of Eminence (IoE), University of Hyderabad (No. UoH/IoE/RC3-21-052), Indian Council of Medical Research (ICMR), GoI (ISRM/12(72)/2020, ID: 2020-2951), and Department of Biotechnology, GoI (No. BUILDER-DBT-BT/INF/22/SP41176/2020) for their financial support. Mallikarjuna T would like to thank ICMR, GoI, for the financial support as SRF (Ref. No: ISRM/11(47)/2019).

## Author contributions

Ayushi Dwivedi: Conceptualization, Methodology, Software, Data curation, Formal analysis, Investigation, Validation, Writing – original draft. Afrin Zulfia S: Methodology, Data curation. Mallikarjuna Thippana: Methodology, Data curation, Investigation. Sai Nikhith Cholleti: Methodology, Data curation, Investigation. Vaibhav Vindal: Conceptualization, Methodology, Software, Resources, Writing – review & editing, Supervision.

